# Natural hypothalamic circuit dynamics underlying object memorization

**DOI:** 10.1101/603936

**Authors:** Christin Kosse, Denis Burdakov

## Abstract

Memorizing encountered objects is fundamental for normal life, but the underlying natural brain activity remains poorly understood. The hypothalamus is historically implicated in memory disorders, but whether and how its endogenous real-time activity affects object memorization remains unknown. We found that upon self-initiated object encounters, hypothalamic melanin-concentrating hormone (MCH) neurons emit dynamic, object-encounter-associated signals encoding object novelty. Optosilencing of these signals, performed in closed-loop with object encounters selectively during object memory acquisition, prevented the ability to recognize the previously encountered objects. Optogenetic and chemogenetic connectivity analyses demonstrated that local GAD65 neurons form an inhibitory GAD65→MCH microcircuit that controls the object-encounter-associated MCH cell signals. GAD65 cell optosilencing during object memory acquisition enhanced future object recognition through MCH-receptor-dependent pathways. These results provide causal evidence that natural, object-associated signals in genetically-distinct but interacting hypothalamic neurons differentially control whether the brain forms object memories.

---

The ability to memorize objects enables one to react differently to novel and previously encountered objects. This ability is fundamental for normal life and impaired in many common and devastating brain pathologies^1, 2^. It is still debated which brain structures and signals are critical for the formation of object memory^3, 4^. Intense research into this topic traditionally focused on brain areas such as the perirhinal cortex and hippocampus^2, 4^. In contrast, causal roles of neural dynamics of the hypothalamus have been underexplored, despite over half a century of evidence implicating this region in memory disorders^5–9^. The hypothalamus contains multiple neuronal types interconnected in complex and poorly-understood ways, including neurons expressing the peptide neurotransmitter melanin-concentrating hormone (MCH)^10^ that innervate many brain areas thought to be important for memory control^8, 11^. While originally MCH_LH_ neurons were only thought to be active during sleep^12^, it was found recently that they are also active during awake spatial exploration^13^. However, it remains unknown whether this natural MCH_LH_ cell activity during wakefulness influences object memory formation, because wakefulness-specific silencing of MCH_LH_ neurons in the context of object memorization has not been performed. It is also unknown what neural circuits shape the MCH_LH_ cell activity during wakefulness, and whether these circuits may control object memory formation. Here we explored these unknowns by temporarily-restricted, reversible silencing MCH_LH_ cells and their upstream neurons (newly discovered here) during object encounters, combined with testing the subsequent behavioral reactions to the previously-encountered vs. novel objects.

Recordings of natural MCH_LH_ cell activity during self-paced navigation in object-containing arenas revealed that MCH_LH_ cells emitted activity bursts when mice encountered objects (Fig. 1A-F, encounter was defined by real-time video-tracking as an entry of mouse nose into the object area, see Fig. 1C and Methods). Such object-related activity was not observed in LH hypocretin/orexin neurons (Fig. S1F), indicating a cell-type-specificity of LH responses to object encounters. When recorded continuously during sequential presentation of novel and familiar objects to the same mice, the novel-object-encounter-associated MCH_LH_ signals decreased as mice spent more time with the object (Fig. 1D-H; Fig. S1E), but increased again when they were presented with a new novel object (Fig. 1D,E; Fig. S1C). When mice were presented with familiar objects, the object-encounter-associated MCH_LH_ signals during consecutive object encounters tended to remain small in amplitude, in contrast to novel-object-associated signals that were initially large and decayed during consecutive object encounters (Fig. 1F-H). This object familiarisation-evoked reduction in MCH_LH_ signals persisted when the familiar object was moved to a new location (Fig. S1B,D), and was maintained for up to 20 h (Fig. S1C,D). Together, these properties of object-encounter-associated MCH_LH_ signals are consistent with signals associated with object memorization.

**Fig. 1.**
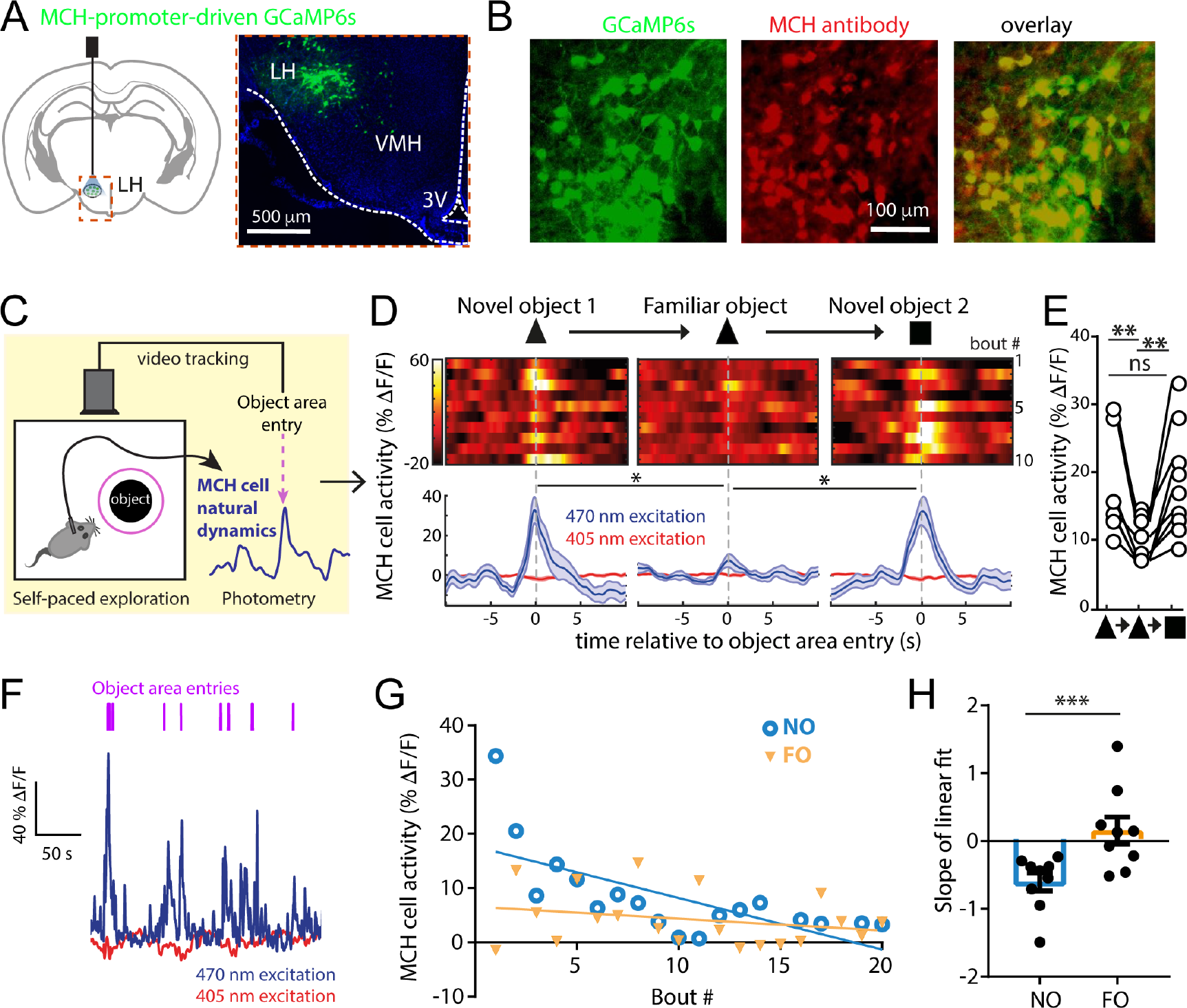
Natural MCH_LH_ cell dynamics underlying sequences of object encounters. **(A)** Targeting scheme (left) and expression (right) of GCaMP6s in MCH_LH_ cells. **(B)** Confirmation of GCaMP6s expression in MCH_LH_ cells (see Methods). **(C)** Schematic (left) of MCH_LH_ cell recording concurrent with behavioral tracking. **(D)** Top, representative heatmaps of MCH_LH_::GCaMP6s fluorescence (at 470 nm excitation) aligned to object-area entry. First 10 object area entries from one mouse (representative data of n = 9 mice). Bottom, group data (means±s.e.m of n = 10 objects visits for one mouse), also showing negative control from 405 nm excitation. One-way RM ANOVA F(2,27)=5.505, p=0.0099, Tukey’s multiple comparisons: Novel object 1(triangle) vs novel object: *p=0.0227, Novel object vs. novel object 2 (rectangle): *p=0.0189 **(E)** Quantification of peak MCH_LH_ cell activity from D, n = 9 mice. One-way RM ANOVA F(1.694, 13.55) = 16.52, p = 0.0004, Tukey’s multiple comparisons post-tests: novel object 1 (triangle symbol) vs novel object 2 (square symbol) p= 0.9991 (ns), novel object 1 vs familiar object (same object presented 30 min later) **p=0.0021, familiar object (middle triangle symbol) vs novel object 2 (square symbol): **p = 0.0073 **(F)** Representative MCH_LH_ cell responses from one mouse to a sequence of self-paced novel object area entries. **(G)** Representative peak MCH_LH_ cell responses from one mouse to a self-paced sequence of novel (NO) and familiar (FO) object area entries; straight lines are linear fits to the data. **(H)** Quantification of data in G for n = 9 mice ***p=0.0074, t(8)=3.556.

To probe whether the object encounter-associated signals of MCH_LH_ cells play a causal role in object memory formation, we close-looped real-time video-tracking of object encounters to MCH_LH_ cell optosilencing in MCH_LH_::ArchT mice (Fig. 2A-C, see Methods). We did this in the context of a classic object memory test ^14, 15^, where mice are exposed to pairs of objects in 2 temporally separated trials (Fig. 2C). In trial 1 (memory acquisition phase) they encountered two identical novel objects, with or without the object-associated MCH_LH_ cell optosilencing (Fig. 2C). In trial 2 (object recognition test), which involved no optosilencing, object memory was quantified as time spent with a novel vs. the previously-encountered object (Fig. 2C, this quantifies object memory since in this test mice are normally less drawn to previously-encountered objects^14, 15^). To prevent variations in sensory or laser exposure from affecting memorization, we tracked the total object encounter time in trial 1, and matched its value across compared conditions (see Methods section “Object recognition tests with controlled familiarization time”). MCH_LH_ cell optosilencing selectively during object encounters in trial 1 prevented MCH::ArchT mice from recognizing the previously-encountered objects in trial 2 (Fig. 2D). In contrast, trial 2 object recognition was normal when the same closed-loop LH laser illumination experiment was performed in control mice lacking the optoinhibitory opsin (MCH_LH_::GCaMP mice) (Fig. 2D). This shows that the natural MCH_LH_ cell activity during initial object encounters is necessary for the object to be treated as familiar in the future, i.e. for object recognition memory formation. The controlled design of these experiments (exposure time matching, mixed order within-mouse repeats, control mice, see Methods) indicates that the disruption of object memory formation by the temporally-targeted optosilencing of MCH_LH_ cells was not due to differences in sensory exposure, or order or laser-related effects.

**Fig. 2.**
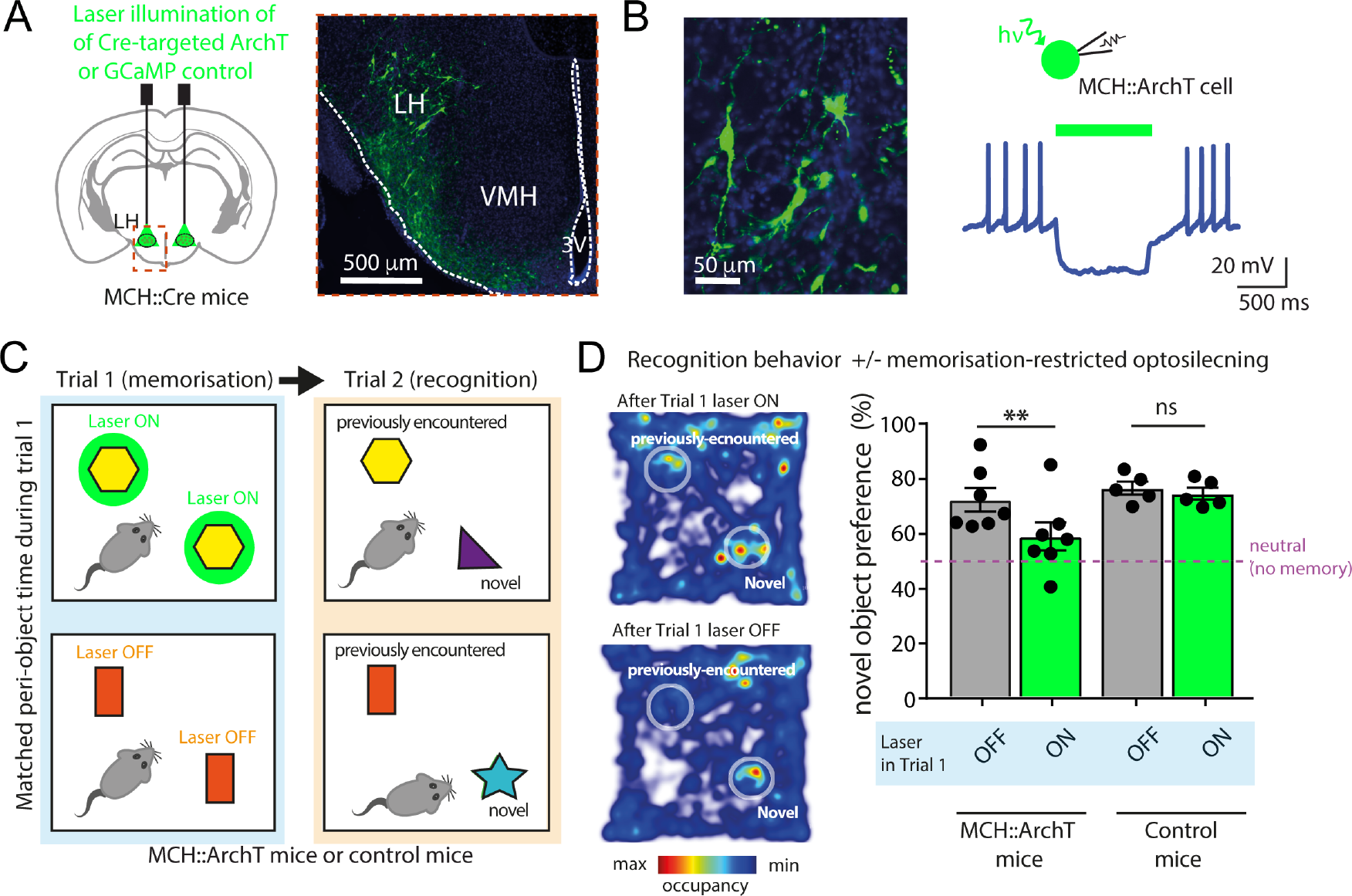
Natural MCH_LH_ cell activity during initial object encounter is required for subsequent object recognition. **(A)** Targeting scheme (left) and expression (right) of ArchT in MCH_LH_ cells. **(B)** Patch-clamp recording (right) confirming silencing of MCH_LH_::ArchT-YFP cells (left) by green light (n = 5 cells). **(C)** Experimental scheme: self-paced exploration of two identical novel objects for the same cumulative peri-object time (trial 1) followed by quantifying exploration of the same arena with one of the previously-explored objects replaced by a novel object (trial 2). **(D)** Left: sample heatmaps showing relative time spent with objects on trial 2. Right: group data. Object-area-entry-coupled bilateral LH laser illumination during trial 1 reduced object recognition in trial 2 in MCH_LH_::ArchT mice (n = 7) but not in control (MCH_LH_::GCaMP) mice (n = 5) (2-way ANOVA: F(1, 8) = 7.43, p=0.0260, Sidak’s multiple comparisons test: **p=0.0034, ns = p = 0.7073). In MCH_LH_::ArchT mice, trial 1 laser ON group, the preference for the novel object during trial 2 was not different from “no memory” (neutral, 50%) criterion (one sample t-test against 50% preference: t(6)=1.775, p = 0.1262), whereas in all other groups significant preference was seen (one sample t-tests against 50% preference: MCH_LH_::ArchT mice laser OFF, t(6)=5.276, p= 0.0019; control mice laser OFF, t(4)=11.37, p = 0.0003; control mice laser ON, t(4)=11.42, p = 0.0003).

The above findings show that inhibition of the object-associated MCH_LH_ activity selectively during memory acquisition is a powerful way to control object memory formation. In search for neural origins of this inhibition, we used channelrhodopsin (ChR2)-assisted circuit mapping to probe functional interactions of MCH_LH_ and neighboring non-MCH cells with local GAD65_LH_ neurons, a recently-characterized LH neural type whose downstream cell targets are yet unknown^16^. In mouse brain slices, optostimulation of GAD65_LH_::ChR2 cells evoked rapid GABAergic inhibitory input in MCH_LH_ cells (Fig. 3A) but not in the neighboring LH orexin/hypocretin cells (Fig. S3B). Optostimulation of MCH_LH_::ChR2 cells did not evoke detectable input in GAD65_LH_ cells (Fig. S3A), suggesting a unidirectional GAD65→MCH LH microcircuit. In complementary *in vivo* circuit-connectivity screens in object-exploring mice, chemogenetic activation of GAD65_LH_::hM3Dq cells (Fig. 3B, see Methods) was able to suppress the novel object encounter-associated activity MCH_LH_ cell bursts (Fig. 3C,D). Thus, a functional inhibitory GAD65→MCH LH circuit exists that is sufficiently powerful to suppress object-encounter-associated MCH_LH_ cell activity.

**Fig. 3.**
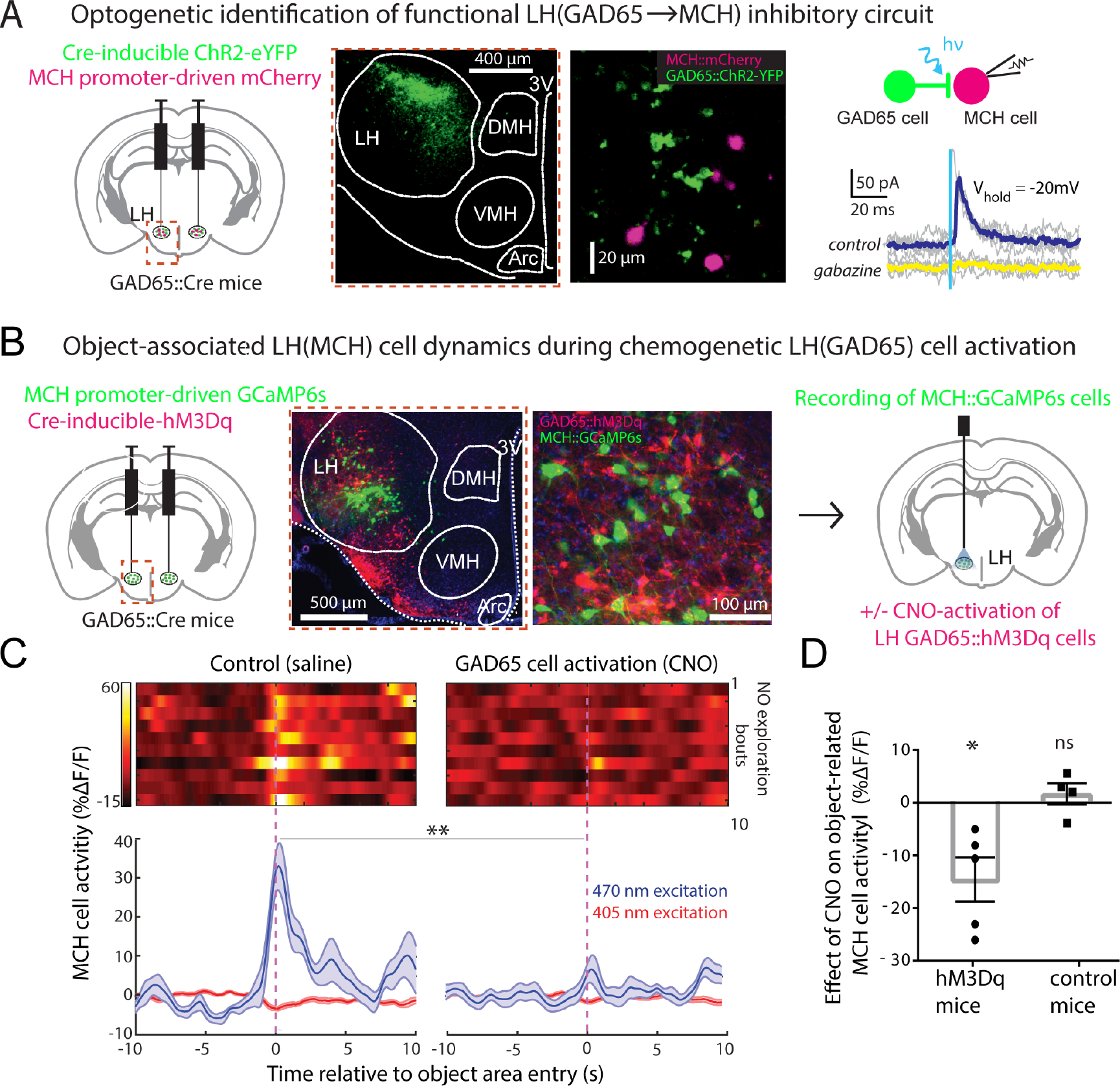
Functional identification of GAD65_LH_ → MCH_LH_ inhibitory circuit. **(A)** Targeting schematic (left) for expression of ChR2 in GAD65_LH_ cells and mCherry in MCH_LH_ cells (middle panels). Right, GAD65_LH_::ChR2 optostimulation evokes a gabazine–sensitive current in MCH_LH_::mCherry cells. Grey lines are individual trials, colored lines are trial averages; n = 14/16 cells were connected; latency between GGA65_LH_ cell optostimulation and the onset of postsynaptic current: 0.83±0.8 ms (n = 14 cells), synaptic current size is quantified in Fig. S3C). (B) Targeting schematic (left) and expression (2 middle panels) of GCaMP6s in MCH_LH_ cells and hM3Dq in GAD65_LH_ cells, for recording of MCH_LH_::GCaMP6s cell activity during GAD65_LH_::hM3Dq cell modulation (right). **(C)** Top, representative heatmaps of MCH_LH_::GCaMP6s fluorescence (at 470 nm excitation) aligned to object-area entry. First 10 object area entries from one mouse (representative data of n = 5 mice). Bottom, corresponding group data (means±s.e.m of n = 10 visits), also showing negative control from 405 nm excitation. Chemogenetic activation of GAD65_LH_::hM3Dq cells decreased object-area-entry-associated MCH_LH_ cell activity peaks (t(18)=3.805, **p= 0.0013, unpaired t-test, representative data comparing 10 object encounters before and after CNO from one mouse, group data are given in D). The late activity at around 5 and 10 s reflect activity outside the object area, which was not investigated further. **(D)** Group data, showing effect of CNO (i.e. response in CNO minus response in saline) on peak peri-object MCH_LH_ cell signals in negative control mice (MCH_LH_::GCaMP6s, n = 4), and in hM3Dq mice (MCH_LH_::GCaMP6s and GAD65_LH_::hM3Dq, n = 5); *p = 0.0257, t(4)=3.466, one-sample t-test, n = 5 mice; ns = p = 0.4623, t(3)=0.8406, one-sample t-test, n = 4 mice.

To investigate whether the natural GAD65_LH_ cell activity influences object memory acquisition via the MCH system, we repeated the memory acquisition –coupled optogenetic interference (Fig. 2) with GAD65_LH_ optosilencing. The GAD65_LH_ cell optosilencing targeted to object encounters during object memory acquisition significantly increased subsequent novel object preference during object recognition (Fig. 4C,D). This indicates that the natural activity of GAD65_LH_ cells opposes object memory formation, as expected from the inhibitory GAD65→MCH LH circuit. If this effect of GAD65_LH_ cells on object memory formation is mediated by the inhibitory GAD65→MCH LH circuit, then it should be diminished by blocking MCH cell outputs. Consistent with this prediction, when the GAD65_LH_ cell optosilencing was performed concurrently with MCH receptor blockade using the MCH receptor antagonist SNAP94847 (20 mg/kg i.p., see Methods), the effect of the GAD65_LH_ cell optosilencing was abolished (Fig 4C,D). Conversely, object memory formation driven by natural MCH signaling (isolated by quantifying behavior with and without SNAP94847 in individual mice) was significantly increased by GAD65_LH_ cell inhibition (Fig. 4E), confirming that MCH_LH_ cells regulate behavior according to GAD65_LH_ cell tone. This shows that the object-encounter-associated, natural GAD65_LH_ cell activity governs object memory formation via MCH-receptor-dependent pathways.

**Fig. 4.**
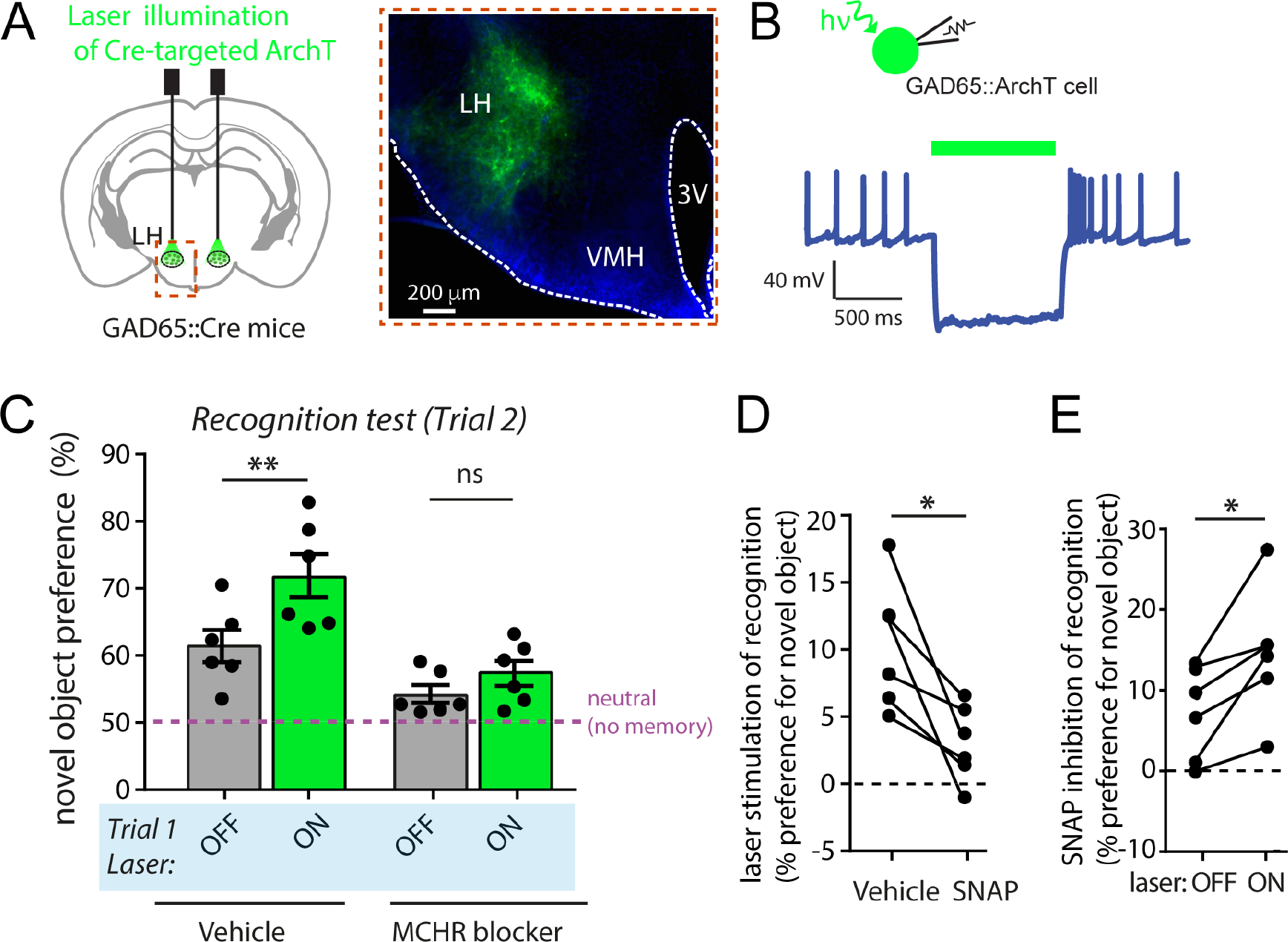
Natural GAD65_LH_ cell activity during initial object encounters influences subsequent object recognition via MCH receptor signaling. **(A)** Targeting schematic (left) and expression (right) of ArchT in GAD65_LH_ cells. (**B**) Patch-clamp recording from GAD65_LH_::ArchT cells confirming cell inhibition by green light (n = 5 cells). (**C**) Role of GAD65_LH_ cell activity in object memory formation (experimental design as in Fig. 2C). Object-area-entry-associated GAD65_LH_ cell optosilencing during Trial 1 increased object recognition in trial 2 in the absence (2-way ANOVA, F(1, 5) = 17.1, p=0.0090, Tukey’s multiple comparisons test: **p = 0.0036) but not presence of MCHR blocker SNAP94847 (2-way ANOVA, F(1, 5) = 17.1, p=0.0090, Tukey’s multiple comparisons test: ns = p = 0.2931), n = 6 mice. (**D**) Quantification of trial 2 object recognition enhancement by the trial 1 GAD65_LH_ cell optosilencing, in the presence and absence of SNAP (n = 6 mice, paired t-test: t(5)=3.488, *p= 0.0175). (E) Quantification of trial 2 object recognition enhancement by the trial 1 SNAP94847, with and without concurrent GAD65_LH_ cell optosilencing, n=6 mice, paired t-test: t(5)=3.488, *p=0.0175.

Overall, these findings show that mice do not recognize previously-encountered objects unless their MCH cells are active during the prior object encounters, and that a novel GAD65→MCH microcircuit governs the size of this MCH cell activation. The type of object recognition memory that we studied is fundamentally important for normal behavior in both humans and animals^2, 17^. To the best of our knowledge, our study is the first to make causal links between object-associated dynamics of hypothalamic neurons and object memory formation. Previous molecular and pharmacological studies of MCH neuropeptide signaling and avoidance memory^8, 9^ did not involve real-time, reversible manipulation of ongoing MCH cell activity at behaviorally-relevant timescales, and thus contained no indication when the natural MCH cell dynamics influences memory, nor how the activity state of specific upstream circuits shapes such memory-gating MCH cell dynamics. The findings presented here therefore make the previously missing causal link between object-associated hypothalamic circuit activity and object recognition memory formation.

These results support broader roles of rapid hypothalamic signals in cognition than previously considered, as also supported by recent data on other hypothalamic neurons ^18–21^. Furthermore, our data suggest that the hypothalamus not only relays a critical input for computation of appropriate behavior, but that local hypothalamic microcircuits also contribute to this computation. This contribution can be critical for fundamentally important behavior, since disrupting natural LH processing transiently and specifically during initial object encounters prevented mice from displaying normal behavioral responses (i.e. recognition) to novel and familiar objects in the future (Figs. 2 and 4). While our study focussed on inanimate objects in order to isolate object memory effects from reward or social motivators, in the future it will be interesting to investigate how local hypothalamic processing affects novelty and familiarity behavior towards more complex objects such as food and conspecifics.

Our finding that GAD65→MCH LH circuit is important for object memory does not rule out that this circuit may also be involved in other functions. While so far we found no evidence that MCH_LH_ or GAD65_LH_ cells are involved in spatial working memory (Fig. S4), nor that MCH cells signal spatial locations (Fig. S1B,D), we did observe they may signal novel sensory qualities of food (Fig. S5). The possibility that this circuit may signal multiple novel sensory experiences does not, in our opinion, undermine the validity and importance of our findings relating to its involvement in object recognition memory. At the neuroanatomical level, MCH_LH_ cell axons and MCH receptors are found brain-wide, including multiple regions speculated to be involved in object memory ^3, 4, 9, 11, 22^, where MCH is proposed to alter synaptic plasticity thus making memories more likely to form ^8, 9^. Probing this broader downstream connectivity of the GAD65→MCH LH circuit will improve our understanding of hypothalamic gating of cognition, though this is likely to be challenging given the breadth of MCH_LH_ projections and the current lack of consensus about relative roles played by different brain regions in object recognition memory. Upstream, it would be interesting to probe whether known regulatory inputs to MCH_LH_ and GAD65_LH_ cells – for example orexin, insulin, and glucose ^13, 16, 23, 24^ – may act to match memory-related processes to stress and energy levels.

In summary, our study identifies a neural circuit that governs brain representations of object novelty, and links the natural object-related activity of this circuit to a vital cognitive function: object recognition memory formation. This previously unknown circuit mechanism for the control of object memory formation offers new insights into neuromodulation of valued cognitive abilities that are key targets of rehabilitation in neuropsychiatric disease.

---

**Fig. S1.**
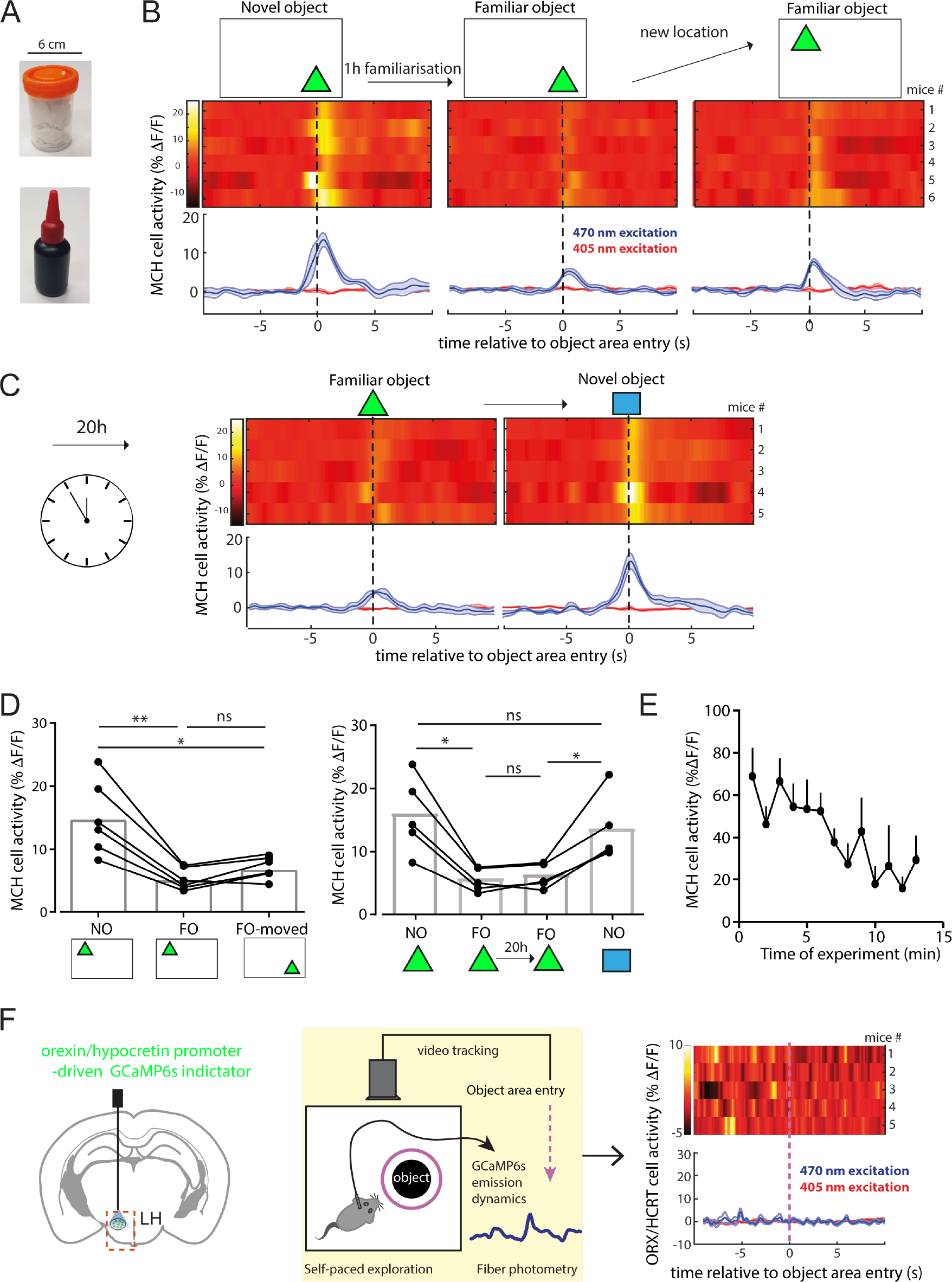
Specificity of MCH_LH_ cells responses to novel objects: roles of object location, memory retention time, and comparison with other LH cells. **(A)** Typical examples of objects used in the study. All objects were approximately the same size but differed in shape. **(B-F)** Control data on hypothalamic representation of object encounters. **(B)**, Moving a familiar object to a new location in the cage does not restore large MCH_LH_ cells responses typical of novel objects, indicating that MCH_LH_ cells selectively represent object novelty rather than object place. Heatmaps of MCH_LH_::GCaMP6s fluorescence aligned to object area entry (each heatmap time-sweep is an average of first 10 entries per mouse; traces below heatmaps are means±s.e.m. of n = 6 mice, also showing negative control fluorescence from 405 nm excitation). **(C)**, Continuation of the experiment shown in B (the experiment was continued in C for 5/6 mice shown in B), showing that the object familiarisation -associated reduction in the MCH_LH_::GCaMP6s signal persists for 20 hours, and can then be reversed by presenting a novel object (n = 5 mice). **(D)**, Left, analysis of group data shown in B: One-way ANOVA F(1.12, 5.601) = 22.05, p = 0.0036, Tuckey’s post test **p = 0.005, *p = 0.0247, ns = p = 0.0863. Right, analysis of group data shown in C: One-way ANOVA F(1.467, 5.867) = 21.38, p = 0.0027, Tuckey’s post test left *p = 0.0186, right *p = 0.0401, ns = p > 0.3… **(E)**, Time-course of novel object area entry-associated MCH cell activity peak size during a typical experiment (means+s.e.m. of n = 5 mice). **(F)** Specificity of MCH cell dynamics: hypocretin/orexin cell activity does not increase during novel object area entry. Left, Calcium indicator targeting (we used a previously validated targeting method characterized in Gonzalez et al, *Current Biology* 2016, 26(18): 2486-2491). Center, Experimental set-up. Right, Hypocretin/orexin_LH_ cell dynamics associated with novel object exploration (data plots are of the type as described above in B, n = 5 mice).

**Fig. S2.**
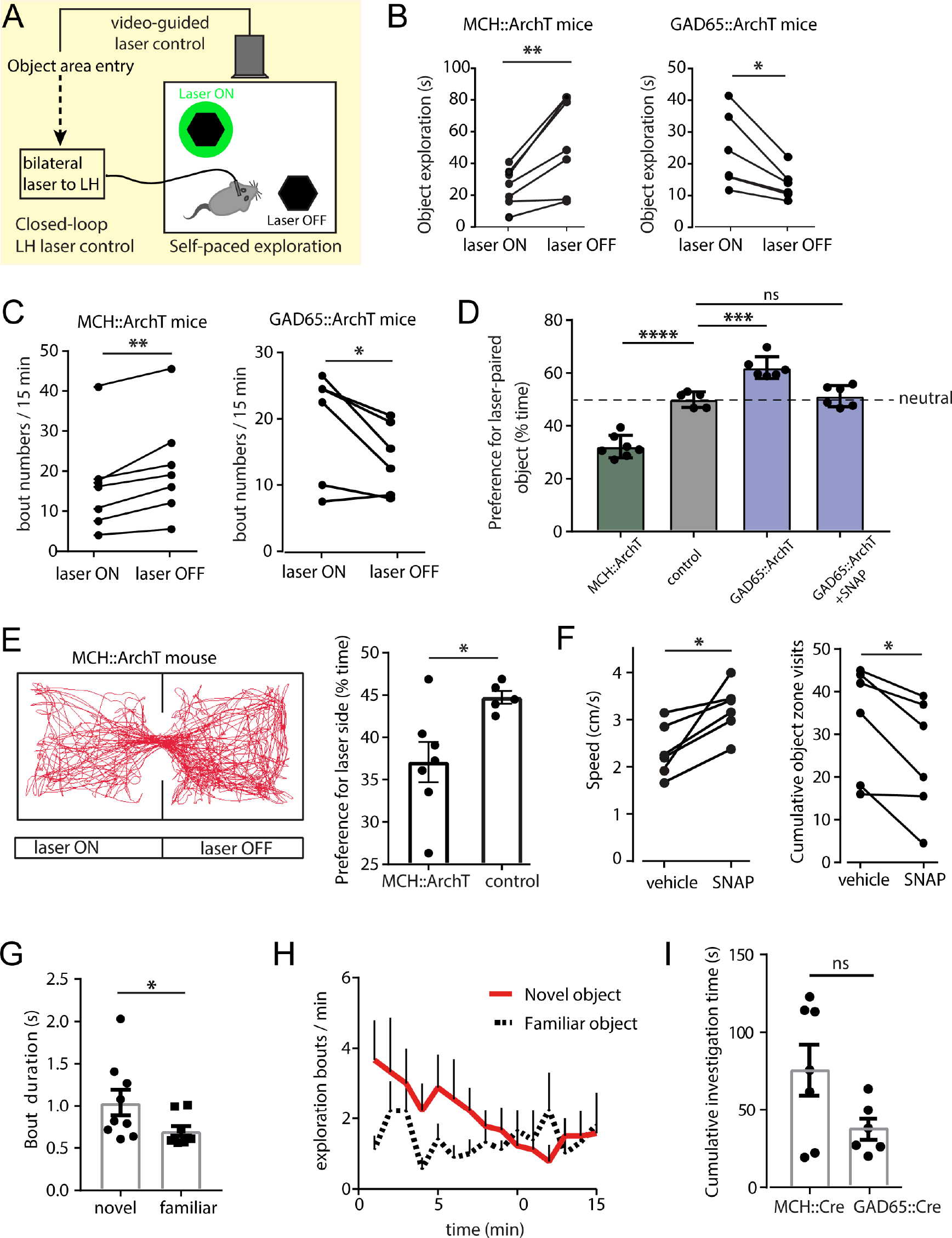
Control experiments that guided our experimental design and interpretation. (**A-D**) Control experiments for effect of optogenetic manipulations on object exploration duration (these effects were subsequently controlled for by cumulative exploration time-matching, as described in Methods). Self-paced exploration of two identical novel objects for 10 min with two identical objects placed diagonally whilst one object and its peri-object space were paired with laser triggering through closed loop video tracking. Object areas were defined as extending by 3 cm around the object, and automated real-time nose video-tracking (Ethovision XP) scored each time the mouse nose entered the object area as a visit. In experiments with MCH receptor blocker, the MCH antagonists SNAP94847 or vehicle were injected i.p. 45 min before the experiment started, and familiar object were familiarised before injection. Objects were assigned according to a crossover design. **(A)** Experimental scheme. **(B)** Quantification of raw data of time spent in peri-object area of the laser ON paired object and control object without laser for MCH::ArchT (left), n = 7 mice, paired t-test: t(6)=3.991, **p= 0.0072 and GAD65::ArchT (right), n=6, paired t-test: t(5)=3.503, *p=0.0172. **(C)**, Quantification of exploration bout numbers at the laser-paired and control peri-object area for MCH::ArchT (left), n = 7 mice, paired t-test: t(6)=4.57,**p=0.0038 and (right) for GAD65::ArchT, n=6, paired t-test: t(5)=2.738, *p= 0.0409. **(D)** Quantification of the relative time spent with the laser-paired peri-object space compared to the overall time of peri-object exploration: One way ANOVA F(3, 20)=65.76, p<0.0001, Dunnett’s multiple comparison test: MCH::ArchT (n=7) vs control (MCH::GCaMP) mice (n=5), ****p=0.0001; control (n=5) vs GAD65::ArchT (n=6), ***p=0.002; control (n=5) vs GAD65::ArchT+SNAP (n=6) ns=p=0.9024. **(E)** Real-time place preference (RTPP) effect of MCH_LH_ cell optosilencing. Left, representative trajectory of a MCH::ArchT mouse in a place preference chamber where one side was paired with bilateral LH laser illumination, showing that the mouse spent more time in the laser off side of the chamber. Right, quantification of the preference (time spent) for the laser on side of the chamber for n=7 MCH::ArchT and n=5 control (LH MCH::GCaMP) mice, unpaired t-test: t(10)=2.607, *p=0.0262. **(F)** Effect of SNAP vs vehicle in LH GAD65::ArchT n=6 mice (left) on speed: paired t-test, t(5)=3.536, *p=0.0166 and (right) on peri-object area visits with two identical novel objects: paired t-test, t(5)=3.731, *p=0.0136. **(G)** Quantification of bout duration (i.e. object area occupancy) during photometry recordings of LH MCH::GCaMP mice (n=9 mice) exploring either novel or familiar objects. Paired t-test: t(8)=2.434, *p=0.0409. **(H)** Quantification of exploration bouts (peri-object area entries) per minute for novel and familiar objects of LH MCH::GCaMP mice (n=9 mice). **(I)** Quantification of overall time spent exploring two identical objects when one is paired with laser illumination for LH GAD65::ArchT (n= 6 mice) and LH MCH::ArchT (n=7 mice), unpaired t-test: t(11)=2.014, ns=p=0.0691.

**Fig. S3.**
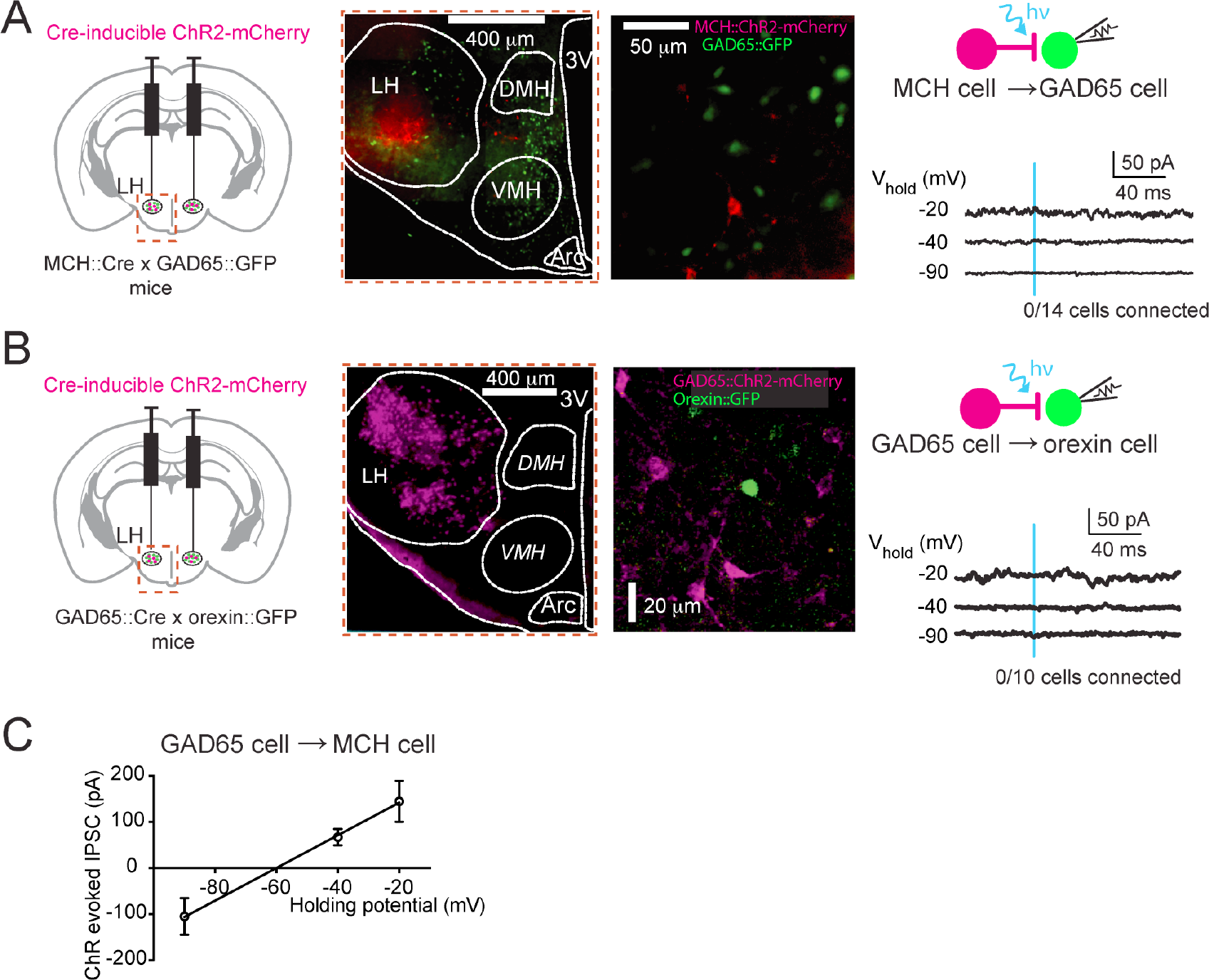
Additional electrophysiological data on LH local connectivity mapping. **(A)** Targeting schematic (left) for expression of ChR2 in MCH_LH_ cells and GFP in GAD65_LH_ cells (middle panels). Right, MCH::ChR2_LH_ optostimulation evokes no currents in GAD65::GFP_LH_ cells (n = 14 cells). **(B)** Targeting schematic (left) for expression of ChR2 in GAD65_LH_ cells and GFP in orexin_LH_ cells (middle panels). Right, GAD65::ChR2_LH_ optostimulation evokes no currents in orexin::GFP_LH_ cells (n = 10 cells). **(C)** Current-voltage relationship of the GAD65_LH_ cell optostimulation-evoked peak inhibitory postsynaptic currents (IPSCs) in MCH_LH_ neurons, at different holding potentials (means±s.e.m. of n=14 cells).

**Fig. S4.**
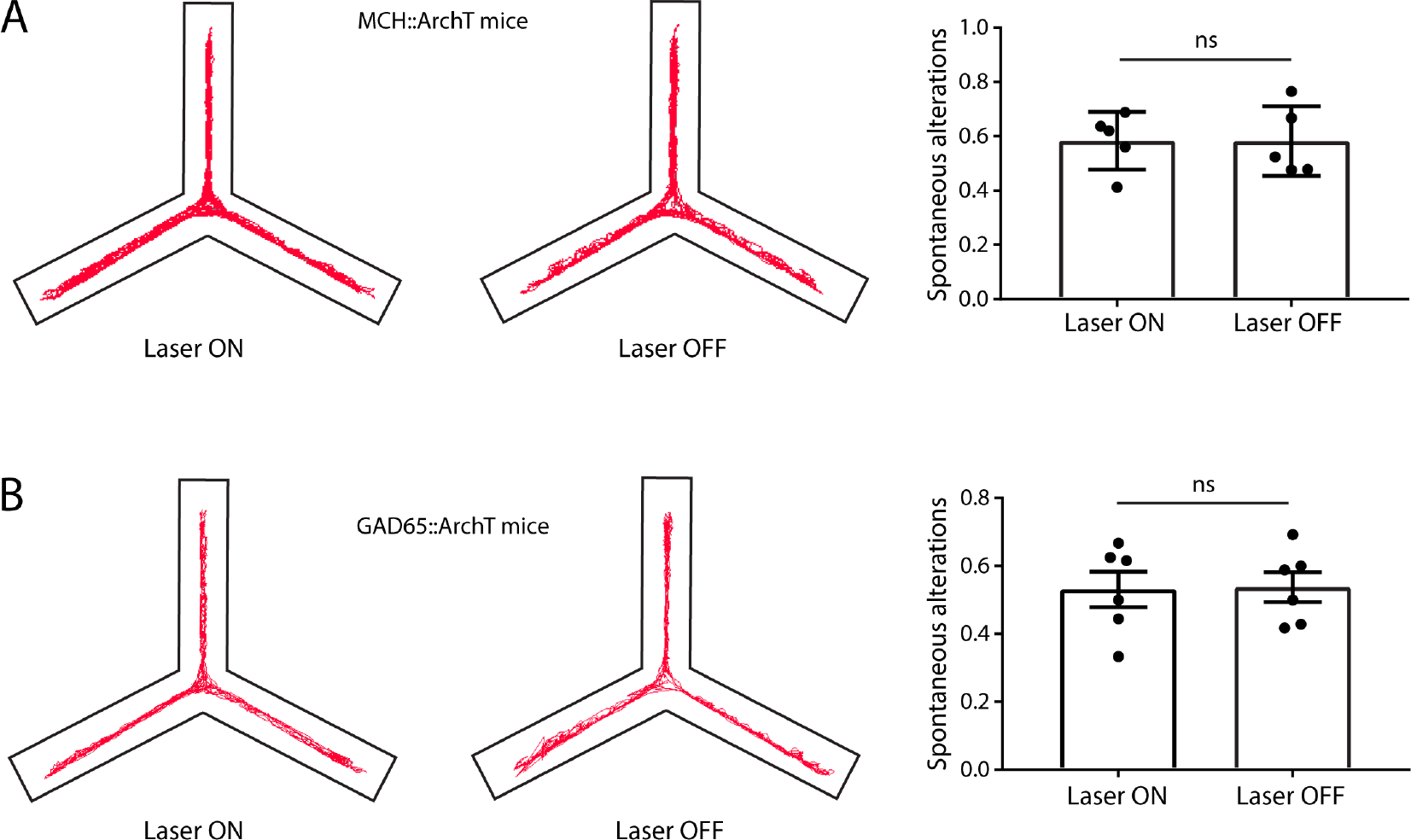
Spontaneous alterations Y maze data. **(A)** Left, representative examples of movement traces of an LH MCH::ArchT mouse in a Y-maze during concurrent laser on and off. Right, group data quantification of the proportion of spontaneous alterations (defined as triad of visits to three different arms), n= 5 MCH::ArchT mice, paired t-test: t(4)=0.02115, ns=p=0.9841. **(B)** Same as (A) but with LH Gad65::ArchT mice, n=6 mice, paired t-test: t(5)=0.172, ns=p=0.8702.

**Fig. S5.**
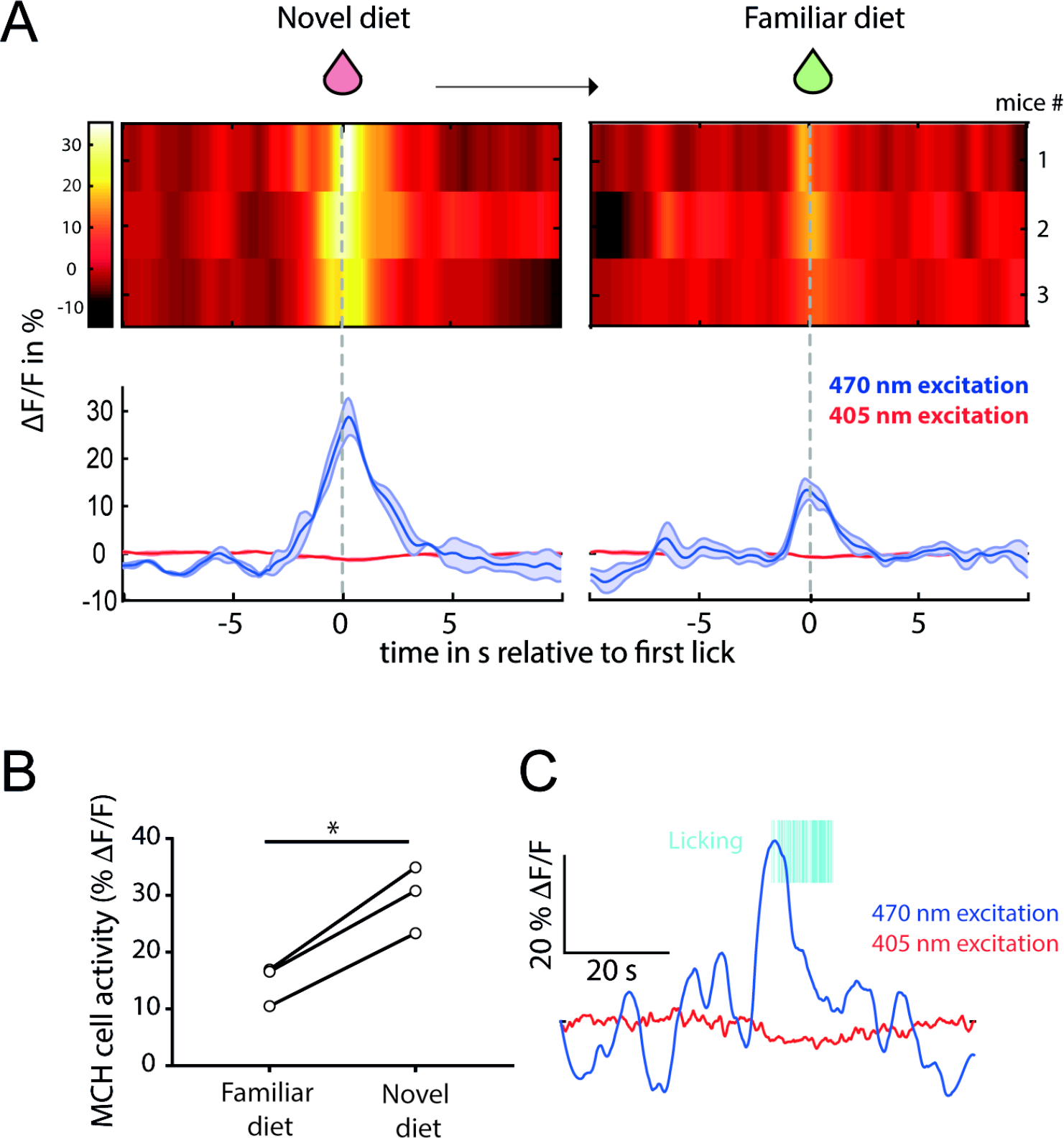
MCH cell responses to consumption of novel and familiar liquid diet. **(A)** Photometry data of n = 3 mice, each an average of recordings aligned to the first lick of the first 3 lick bouts in response to novel and familiar liquid diets (strawberry milkshake or apple juice). Licks were recorded and times-tamped with a lick sensor connected to the food spout (method described in Gonzalez et al, Current Biology 2016, 26: 2486-2491). Heatmaps represent the averaged data of one mouse per line, whist the graph below shows an average of the heatmap data. **(B)** Quantification of data in **(A)** comparing the peak activity of n = 3 mice, paired t-test: t(2)=9.727, *p=0.0104. **(C)** Representative example of raw data showing the photometry data during 405nm and 470nm excitation and simultaneous touch sensor recordings of the spout delivering the novel liquid diet.

## Acknowledgements

The experiments in this study were supported by the Francis Crick Institute (grant code FC001055), which receives its core funding from the UK Medical Research Council, the Wellcome Trust, and Cancer Research UK. CK designed the study, carried out the experiments, and co-wrote the text. DB conceived the study, supervised the experiments, and wrote the text. We thank Dr. Daria Peleg-Raibstein and Cristina Concetti for critical reading of the manuscript. The authors have no competing interests. All data are available in the manuscript of the supplementary materials.

## METHODS

### Genetic targeting

All procedures followed United Kingdom Home Office regulations and were approved by the Animal Welfare and Ethical Review Panel of the Francis Crick Institute. Mice were kept on a standard 12-h/12-h light/dark cycle and on standard mouse chow and water ad libitum. Adult male and female mice (at least 8 wk old) were used for in vitro experiments. Adult male mice were used for behavioral experiments, which were performed during the dark phase. The following previously characterized and validated transgenic mouse lines (or their crosses) were used, where indicated: MCH::Cre mice ^1^, GAD65::Cre mice ^2^, GAD65::GFP mice ^3^, orexin::GFP mice ^4^. The GAD65::Cre mice were bred in homozygous (hom)-WT pairs with C57BL/6 mice; all other transgenic mice were bred in het-WT pairs with C57BL/6 mice. For brain surgeries, mice were anesthetized with isoflurane and injected with meloxicam (2 mg/kg of body weight, s.c.) for analgesia. After placement into a stereotaxic frame (David Kopf Instruments), a craniotomy was performed and a borosilicate glass pipette was used to inject viral vectors bilaterally into the LH. Two injections (each 75 nL) were made into the LH in each hemisphere (bregma: −1.30 mm, midline: ±1 mm, from brain surface: 5.20 mm and 5.25 mm). Before any manipulations, mice were allowed to recover from surgery for at least 1 wk after surgery while single-housed. To target expression of the activity indicator GCaMP6s to MCH_LH_ neurons, we used an AAV vector carrying the 0.9kb preproMCH gene promoter ^5^, AAV9.pMCH.GCaMP6s.hGH (1.78×10^14 gc/mL; Vigene Biosciences). The specificity of GCaMP6s expression was confirmed by staining with MCH antibody (Fig. 1B, 93% specificity was observed by analysis of MCH immunoreactivity colocalisation in 426 GCaMP6s neurons from 3 brains). For optogenetic silencing of MCH_LH_ or GAD65_LH_ neurons, we injected Cre-dependent AAV8.Flex-ArchT-GFP (4.6×10^12 gc/ml; UNC Vector Core) into LH of the MCH::Cre or GAD65::Cre mice, respectively. For ChR-assisted circuit mapping, “FLEX switch” ChR2 constructs were injected into LH of the MCH::Cre or GAD65::Cre mice, as indicated. These constructs were either AAV1.EF1.flox.hChR2(H134R)-mCherry.WPRE.hGH (8.78 × 10^12 gc/mL; UPenn Vector Core) or AAV1.EF1.DIO.hChR2(H134R)-YFP.WPRE.hGH (6.2 × 10^12 gc/mL; UPenn Vector Core). Cre-dependent “DREADD” chemogenetic actuator hM3Dq was targeted to GAD65_LH_ neurons in GAD65::Cre mice by injecting the vector AAV8.hSvn-DIO-hm3D(Gq)-mCherry (2.2 × 1012 genome copies (gc)/mL; UNC Vector Core) into LH of the GAD65::Cre mice, as described and functionally validated by demonstration of CNO-induced inhibition of GAD65::hM3Dq neurons in our previous work ^6^.

### Fiber photometry

After LH injection of MCH-promoter-driven GCaMP6s, alone in C57/Bl6 mice or in combination with the Cre-dependent activatory DREADD (hM3Dq) in GAD65::Cre mice, fiberoptic implants were stereotaxically installed with the fiber tip above the LH (1.35 mm caudal from bregma, 1.0 mm lateral from midline, and 5 mm ventral from brain surface) and fixed to the skull as in our previous work ^7 8^. This method is estimated to capture fluorescence signals from within ≈ 500 μm of the fiber tip ^7^. Fiber tip locations were verified in each mouse by examining slices with a visible fiber tract. Fiber photometry was performed as described ^7^, but the excitation model was modified to provide interleaved 405 nm and 470 nm excitation light pulses via LEDs ^9^. Fluorescence emission produced by 405 nm excitation is not sensitive to calcium and thus provides a real-time control for motion artefacts ^9^. Fluorescence signals were normalised to produce the plotted % ΔF/F values as follows: ΔF/F = 100 * (Fr − F)/F, where Fr is the raw signal and F is the mean of the first 10s of trial. Before photometry recordings, mice were habituated to the recording chamber, the plugging in procedure, and (where relevant) i.p. injections. On the day of fibre photometry recordings, mice were given 10 min to adjust to the chamber before an object was introduced. During the next 1h mice had time to familiarise themselves with the object, after which the object was removed (i.p. injections of CNO or saline where performed where relevant at this point). 30 min after this, the familiar object was reintroduced and mice had free access to explore it while their brain signals were recorded, and head location was video-tracked (Ethovision XP, 15 frames/s). The novel object trial followed, by exchanging the familiar object for a novel object and allowing mice to freely explore the novel object while their brain signals were recorded, and head location was video-tracked. Object exploration bouts were detected by nose video-tracking (Ethovision XP), and their onset defined as the first frame when the mouse nose entered the object area (defined as a 3 cm –wide perimeter around the object). Choice of objects for familiar and novel trials were based on a crossover design to avoid any confounding factors due to differences in objects. To compare MCH::GCaMP6s_LH_ calcium signals between mice, we selected the first 10 exploration bouts of each mouse for the novel and familiar objects, and used these data to derive averaged signal per mouse. During the first 10 entries the object was investigated more frequently if it was novel (Fig. S2H), and since our aim was to define neural correlates of behavioral responses to novel objects, we chose the first 10 entries for analysis of MCH photometry signals.

### Cell-type-specific optosilencing close-looped to object encounters

Mice were bilaterally LH-injected with Cre-dependent ArchT (or, in control experiments, Cre-dependent GCaMP6s), and bilaterally implanted with intra-LH optical fibres using the coordinates and procedures as described above for fiber photometry. 3 weeks after surgery, mice were handled and habituated to the recording arena before any procedure started. For experiments, a green laser (532 nm, LaserGlow) was connected to the bilateral fibre implants to yield ≈20 mW light power output at the fiber tip. Since photometry recordings showed an onset of increased MCH neuron activity before mice entered the object area (Fig. 1D), we paired bilateral LH laser illumination with times when mouse nose was <2 cm away from object area (i.e. within 5 cm perimeter from object). For control experiments investigating the effect of silencing GAD65_LH_ or MCH_LH_ neurons on object exploration, mice were freely behaving for 10 min in an open field arena with two identical objects, and the peri-object area of one object (defined as above) was paired with the bilateral LH laser illumination to test for the GAD65_LH_ or MCH_LH_ neuron effects on object exploration (Fig. S2A-D). The propensities for self-paced object investigation of GAD65::Cre and MCH::Cre mice were investigated in control experiments and found to be similar (Fig. S2I).

### Object recognition tests with controlled familiarisation time

For object recognition memory tests^10^ (Fig. 2C,D; Fig. 4C-E), during the laser ON familiarisation, the bilateral LH laser illumination was triggered whenever the mouse entered the peri-object area (as defined above) of either object. No laser was applied during recognition trials. Laser OFF familiarisation was performed in the same mice with the same temporal contingencies as laser ON familiarisation, but with a new set of objects. After 1 h of retention interval, during which mice were returned to their home cages and no experimental manipulations were performed, the recognition trial (=second trial) consisted of 10 min during which mice freely explored one object from the previous familiarisation trial (familiar object) and one novel object. Sets of novel and familiar objects were alternated between mice in a crossover design. For novel object recognition tests where MCH receptors were blocked with SNAP 94847 (Fig. 4C-E), mice were i.p. injected with SNAP or vehicle solutions 45 min before trial 1. In these experiments, a longer interval between familiarisation/acquisition and recognition trials was used (20 h), to ensure that MCH receptors were only blocked during memory acquisition, and that mice were unimpaired by SNAP during recognition tests.

Our aim was to specifically examine the effects of LH optosilencing on memory formation, independently of factors such as the duration of sensory exposure to objects during familiarisation/memorisation. Therefore, in laser ON and laser OFF familiarization trials, a constant cumulative exposure of mice to objects was imposed, by real-time videotracking the cumulative object encounter time (time when the mouse nose was in the object area), and terminating all trials when the same cumulative object encounter time (30 s) was reached. This ensured that differences in object memory acquisition were not due to variation in initial object exposure between different mice or trials, or different optosilencing conditions that may otherwise have influenced the total object investigation time as suggested by our control experiments (Fig. S2A-D).

### Y maze test of spatial memory

Continuous spontaneous alterations in a Y maze were measured with and without concurrent optogenetic silencing in the same mice (sequence of optogenetic silencing and laser off was alternated between mice) (Fig. S4). Mice were connected to bilateral patch cords 10min before start of the experiments and then transferred to the centre of a standard Y maze (3 arms, 30cm long, 120º apart). During the following 8 min, mice were free to explore the arms of the Y maze whilst video tracking with Noldus Ethovision scored the spontaneous alterations defined as consecutive entries into three different arms ^11^ ^12^ ^11^.

### Experimental sequences in behavioral experiments

Crossover-like experimental designs were used in all *in vivo* photometry and optogenetic experiments, to prevent artefacts and biases and isolate the effects of variables under investigation. Specifically, presentations of novel and familiar object were alternated within and between mice to avoid behavioral fatigue or order effects. Photometry experiments were designed to expose the same mouse to sequences of novel and familiar objects that avoided behavioral habituation or calcium indicator degradation as confounding factors (e.g. novel→familiar→novel, Fig. 1D, Fig. S1B,C). Optogenetic experiments were based on a crossover-like design where manipulations involving drugs, laser light, or mouse genotype were arranged in a Latin square to avoid any confounding factors due to day to day differences or carry-over effects. To prevent potential arena side biases from influencing the results of experiments involving 2 objects positioned at different sides of arena, trials were repeated with laser OFF and ON sides reversed; the presented results are an average of both trials.

### Channelrhodopsin-assisted circuit mapping in brain slices

Brain slice patch-clamp recordings combined with optogenetics were carried out as described in detail in our previous work ^6, 13^. Briefly, LH slices were prepared at least 2 months after virus injection. Coronal brain slices of 250-μm thickness containing the LH were cut while immersed in ice-cold slicing solution. Slices were incubated for 1 h in artificial cerebrospinal fluid (ACSF) at 35 °C, and then transferred to a submerged-type recording chamber. Neurons containing fluorescent markers were visualized with an Olympus BX61WI microscope with an oblique condenser and fluorescence filters. Excitation light was delivered from a LAMBDA DG-5 beam switcher (Sutter) with a xenon lamp and ET470/40 (for ChR2) or ET500/20 (for ArchT) bandpass filters. A 40X 0.8NA objective was used to deliver pulses of excitation light (∼10 mW/mm2, 1 ms for ChR2 activation, or 1 s for ArchT activation) around the recorded cell, and postsynaptic responses were recorded in voltage-clamp (for circuit mapping) or current-clamp (for confirmation of ArchT-mediated photinhibition). Functional ChR2 expression was confirmed by recording light-activated action potentials in the target cells (n = 3 cells per group, not shown). For testing LH output connections of GAD65_LH_ cells, we chose LH neurons based on their genetic markers (MCH::GFP, orexin::GFP, GAD65::GFP) without noting GAD65 fiber location. However, GAD65 fibers were dense and abundant everywhere in the LH ^6^.

### Chemicals and Solutions

For brain slice recordings, ACSF and ice-cold slicing solution were gassed with 95% O2 and 5% CO2, and contained the following: 125 mM NaCl ACSF, 2.5 mM KCl, 1 mM MgCl2, 2 mM CaCl2, 1.2 mM NaH2PO4, 21 mM NaHCO3, 2 mM D-(+)-glucose, 0.1 mM Na+-pyruvate, and 0.4 mM ascorbic acid. The slicing solution contained 2.5 mM KCl, 1.3 mM NaH2PO⋅H2O, 26.0 mM NaHCO3, 213.3 mM sucrose, 10.0 mM D-(+)-glucose, 2.0 mM MgCl2, and 2.0 mM CaCl2. For standard whole-cell recordings, pipettes were filled with intracellular solution containing the following: 120 mM K-gluconate, 10 mM KCl, 10 mM Hepes, 0.1 mM EGTA, 4 mM K2ATP, 2 mM Na2ATP, 0.3 mM Na2GTP, and 2 mM MgCl2 (pH 7.3) with KOH. Gabazine (3 μm) was used where indicated. For *in vivo* chemogenetic manipulations, CNO was injected i.p. at 0.5 mg/kg body weight in experiments involving hM3Dq. The MCH receptor antagonist SNAP 94847 hydrochloride was injected i.p. at 20mg/kg body weight (based on ^14^) after being dissolved in distilled water with 10% DMSO and 30mg/ml (2-Hydroxypropyl)-β-cyclodextrin. All chemicals were from Sigma or Tocris Bioscience.

### Immunohistochemistry

For the immunolabeling of MCH neurons, 50-μm cryosections of pMCH-dependent GCaMP6s injected C57B/l6 mice were stained for MCH with a rabbit antibody to MCH (H-070-47,1:2000, Phoenix Pharmaceuticals) as a primary antibody, and Alexa 555–conjugated donkey antibody to rabbit IgG (A-21244, 1:500, Invitrogen) as a secondary antibody. Slices were then imaged with an Olympus VS120 slide scanner microscope and double labelling of GCaMP with Alexa 555 was quantified with ImageJ.

### Statistical Analyses

Statistical tests and descriptive statistics were performed as specified in the figure legends. All experimental animals were included in the analyses (no pre-selection or exclusion). In each experimental dataset at the cellular level, each n was a different cell (no repeated trials from the same cell were used as n values) and cells from at least three mice were analyzed. Before performing parametric tests, data were assessed for normality with a D’Agostino–Pearson omnibus test or Kolmogorov–Smirnov test for small sample sizes. To compare interactions within normally distributed data with repeated measurements, repeated measures ANOVA was used, with multiple comparison tests where appropriate. All statistical tests are two tailed unless otherwise stated. All error bars indicate the standard error of the mean. Analysis was performed with GraphPad Prism and MATLAB (The MathWorks, Inc.).

### Data and code availability

The datasets generated during and analysed during the current study are available from the corresponding author on reasonable request. Custom codes for the acquisition of photometry data (Labview) and for the analysis of photometry data (Matlab) are available from the corresponding author on reasonable request.

